# Step-by-Step Maturation Mechanism of Binary Toxin Pore Revealed by Cryo-EM Analysis

**DOI:** 10.1101/2024.10.10.617581

**Authors:** Tomohito Yamada, Yukihiko Sugita, Toru Yoshida, Takeshi Noda, Hideaki Tsuge

## Abstract

Membrane pore-forming proteins (PFPs) form ring-shaped membrane-translocating oligomers on membranes, contributing to infection, immunity, and cell death functions. Binary toxins produced by some bacteria consist of an enzymatic component that acts as a toxin and a membrane-binding component that forms a pore that delivers the enzymatic component into target cells. Cryo-electron microscopy (cryo-EM) has advanced our understanding of these translocation mechanisms by revealing several binary toxin complexes’ structures. However, the mechanisms underlying the initial pore formation remain unclear. We determined the structures of several oligomeric forms of the membrane-binding component Ib of the Iota toxin from *Clostridium perfringens* at various stages of pore formation. Structural comparisons revealed how the symmetrically arranged soluble oligomer (prepore) asymmetrically mature into a transmembrane oligomer (pore). These findings enhance our understanding of mechanisms of PFP and provide a structural basis for developing nanodevices using membrane pores.

**Significance Statement:** Although the mode of action of pore-forming proteins (PFPs) has been increasingly elucidated, the structural mechanisms underlying their maturation into membrane-spanning pores remain elusive. One of the key challenges is the difficulty in capturing the intermediate states between the soluble prepore and the mature pore, and its structural analysis. In this study, we successfully prepared the super-complex formed by a bacterial toxin PFP. The super-complex inhibited the constituting oligomers to maturate into the pore, which enabled structural determination of multiple intermediate state. Comparative structural analysis revealed that the transition from prepore to pore occurs through stepwise, domino-like conformational changes. The analysis workflow and insights from this study are expected to advance the field of PFP research.

---

Membrane pore-forming proteins (PFPs) are a family of proteins produced by many organisms, including bacteria, which form homogeneous or heterogeneous multimeric pores on target biological membranes. These proteins involve various vital functions, including cell death, immunity, and infection^1,2^. PFPs are produced as soluble monomers that form ring-shaped oligomers and subsequently create membrane-spanning α-helices or β-barrels to form pores in various biological membranes. Especially, toxic PFPs are known as pore-forming toxins (PFTs). These PFTs can disrupt the homeostasis of target cells by forming pores, leading to the efflux of ions, amino acids, and water and ultimately inducing cell death.

*Clostridium perfringens*^3^, *Clostridioides difficile*^4^, *Clostridium spiroforme*^5^, *Clostridium botulinum*^6^ and *Bacillus anthracis*^7^ produce binary toxins (Iota toxin; Ia and Ib, CDT; CDTa and CDTb, CST; CSTa and CSTb, C2 toxin; C2I and C2II, Anthrax toxin; LF, EF, and PA) composed of enzymatic component and membrane-binding component which function as PFPs^8^. Binary toxins are classified into two groups: intestinal toxins from Clostridium, including the Iota toxin, CDT, CST, and C2 toxin, which have homologous enzymatic components such as actin ADP-ribosyltransferase. The other is the anthrax toxin, whose enzyme components (EF and LF) differ from those of clostridial ADP-ribosyltransferases in both structure and function. However, the membrane-binding components of these toxins are highly conserved, except for their receptor-binding domains. The addition of the enzymatic component alone to cells or organisms does not result in activity, whereas the membrane-binding component alone induces necrosis and dermonecrosis^9,10^. However, components A and B co-addition increases toxicity compared to their individual addition, inducing morphological changes and cell death^11^.

Cell entry mechanisms are similar for clostridial binary toxins and anthrax toxins. The monomeric membrane-binding component binds to the target cell receptor and is activated by proteases to form a soluble ring-shaped oligomer (prepore). The endosomes take up the enzymatic components and prepores. As the endosome acidifies, the pore undergoes a conformational change into a membrane-spanning β-barrel pore, allowing the enzymatic component to translocate into the cell^12,13^. The translocated enzymatic component exerts severe effects on host cells through actin depolymerisation via ADP-ribosylation activity by clostridial binary toxins^6,14,15^, or inhibition of signalling pathways through degradation of MAPKK in anthrax toxin^16,17^. At first, it was shown that the pore of anthrax toxin PA forms the narrowest constriction site (ϕ-clamp) with a diameter of only 6 Å in the centre^18,19^. Understanding how the enzymatic component passes through the narrow ϕ-clamp is key interest in studying membrane translocation mechanisms of binary toxins. We previously determined the cryo-electron microscopy (cryo-EM) structures of the enzyme-pore complexes of the iota and CDT toxins^20,21^. The constriction sites of Ib and CDTb, as named ϕ-clamp in PA, were conserved among Ib and CDTb. We found that after complex formation with Ca-edge and NSQ-loop, which are constriction-sites of the pore, the N-terminal α-helix of the enzymatic component tilts towards the ϕ-clamp, and this structure further unfolds^20,21^. In the case of anthrax toxin, the first α-helix of LF and EF unfold and dock into a deep amphipathic cleft called the α-clamp^22^. Thus, N-terminal unfolding is commonly observed in clostridial binary toxins and anthrax toxins; however, they use different binding modes. It is hypothesised that the disordered structure passes through the ϕ-clamp and refolds its secondary structure within the β-barrel or cytosol. As described above, binary toxin membrane pores are not only pore-forming toxins but also serve as an intriguing research subject because of their unique translocation mechanism involving the unfolding of the enzymatic component.

Mutations in the F427 residue forming the ϕ-clamp of the anthrax toxin PA-pore, as well as the surrounding residues, have significant effects on the maturation from prepore to pore and the translocation activity of the enzymatic component^23–25^. In anthrax PA, point mutation of the D425A which is located at two residues upstream of F427, does not affect the oligomerisation ability, but inhibits the conformational change to the pore state. Cryo-EM structural analysis of this mutant revealed that the oligomer was formed by PA protomers in multiple intermediate states between prepore and pore conformations^26^. Furthermore, adjacent protomers have similar structural states, suggesting that maturation from prepores to pores proceeds in a domino-like manner within the heptamer. However, the complete heptamer structure of each intermediate in this domino-like pore maturation process has not yet been experimentally revealed.

By chance, we obtained a supercomplex in which multiple Ib oligomers were assembled radially (Ib-rosette) while preparing the wild-type Ib component of the iota toxin with liposome. A single particle structural analysis of the Ib-rosette sample revealed that it consisted of both Ib-pores with formed β-barrels and Ib-prepores without the β-barrels. A detailed analysis of the prepore class allowed us to determine the structure of a seven-fold symmetric prepore for the first time in Ib, and multiple asymmetric prepore structures. Comparison with previously reported structures revealed that the protomers forming these prepores were in an intermediate state between the prepore and pore states. Furthermore, comparison of the series of obtained overall structures suggested that the symmetrically arranged protomers underwent a partial loss of symmetry, and this structural change proceeded step-by-step in a domino-like manner to mature into pores. In this paper, we report on the pore maturation process deduced from comparing multiple prepore structures, and high-resolution pore structure obtained in this study.

## Results

### Cryo-EM single particle analysis of Ib-rosette

While attempting to prepare Ib-pores reconstituted in liposomes, the obtained sample was observed using negative staining and cryo-EM revealing that the structure consisted of multiple radially assembled Ib oligomers (referred to as Ib-rosette) **(Extended Data Fig. 1a, b)**. Focusing on the units of the oligomers, single particle structural analysis using CryoSPARC^27^ revealed that Ib-rosette contained not only particles of Ib-pores with membrane-spanning regions covered by lipid micelles, but also particles of Ib-prepores where the β-barrel was not yet formed (**Extended Data Fig. 1c-g)**. The prepore particle data set could be further classified into five subclasses: particles with an open central (class 1) region and particles with a constricted central structure where the base of the β-barrel has formed but is still immature (class 5) (**Extended Data Fig. 2a)**. Class 2−4 suggested that the particle data set was composed of protomers in multiple states, because the density of some portion in the protomers were averaged out. Detailed structural analysis at the protomer level with symmetry expansion (**Extended Data Fig. 2b)**, focused 3D classification (**Extended Data Fig. 2c-d)**, refinement using appropriate reference map for each class and further focused 3D classification (**Extended Data Fig. 2e-g)**, revealed that the heptameric prepore state is formed by three distinct protomer states, which differ from those in the pore state: the initial state, early intermediate state (early int.), and late intermediate (late int.). Finally, the particle data sets for prepore class 1−4 were classified into eight classes, reflecting the different proportions of these three states **(Fig. 1a-j, Extended Data Fig. 2h-i).** The structural characteristics of each protomer’s state are described in the following sections **(Fig. 2)**.

**Figure 1.**
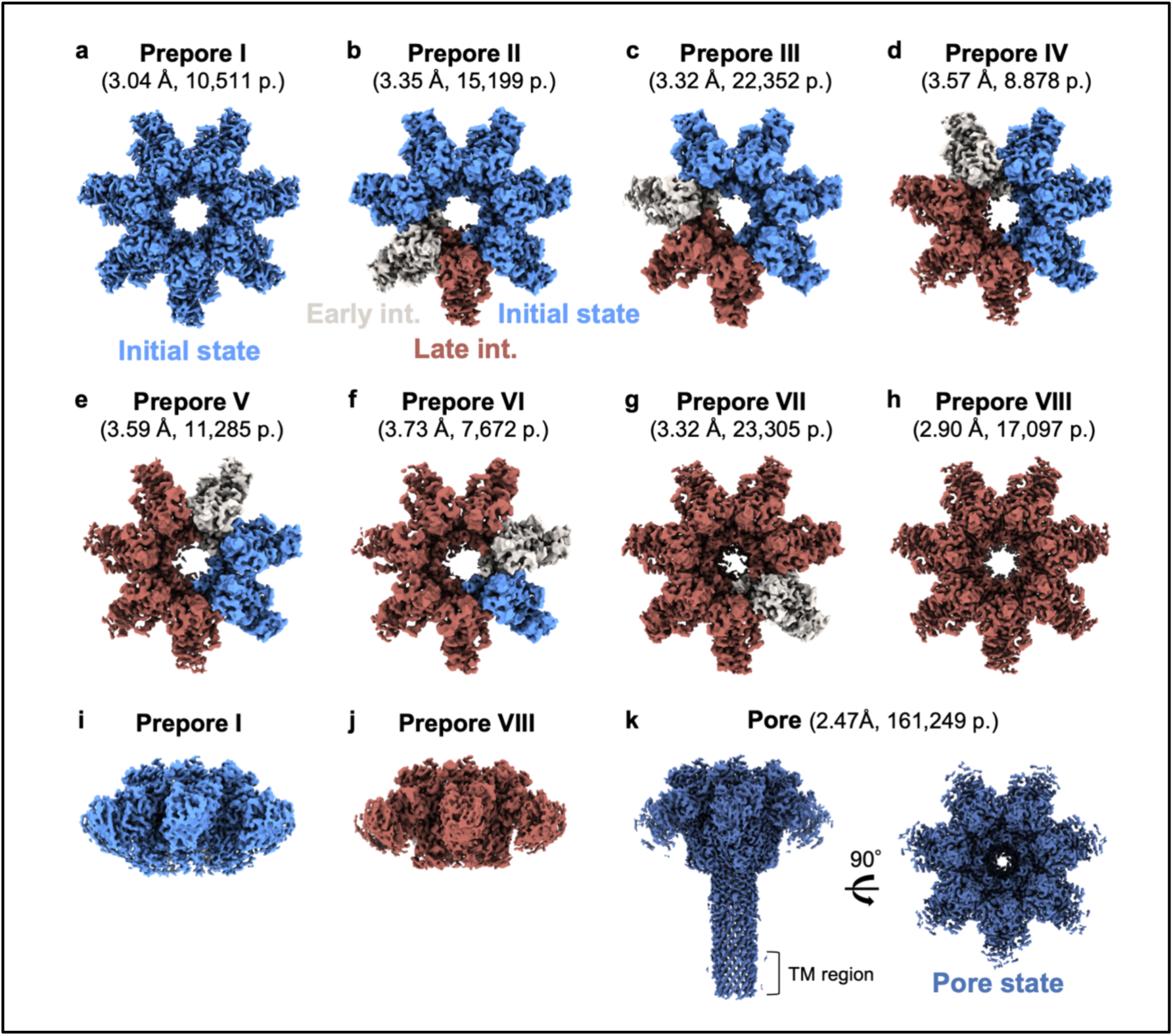
Cryo-EM maps obtained from Ib-rosette analysis. **a−j** Cryo-EM maps of Ib-prepores obtained from cryo-EM single-particle analysis. **a−h** are views from the top of the prepore where the enzymatic component Ia docks. **i, j** Side views of Prepore I and Prepore VIII indicate that the membrane-spanning-barrel has not yet formed at these stages. **k** Cryo-EM map of Ib-pore viewed from the side (left) and top (right).

**Figure 2.**
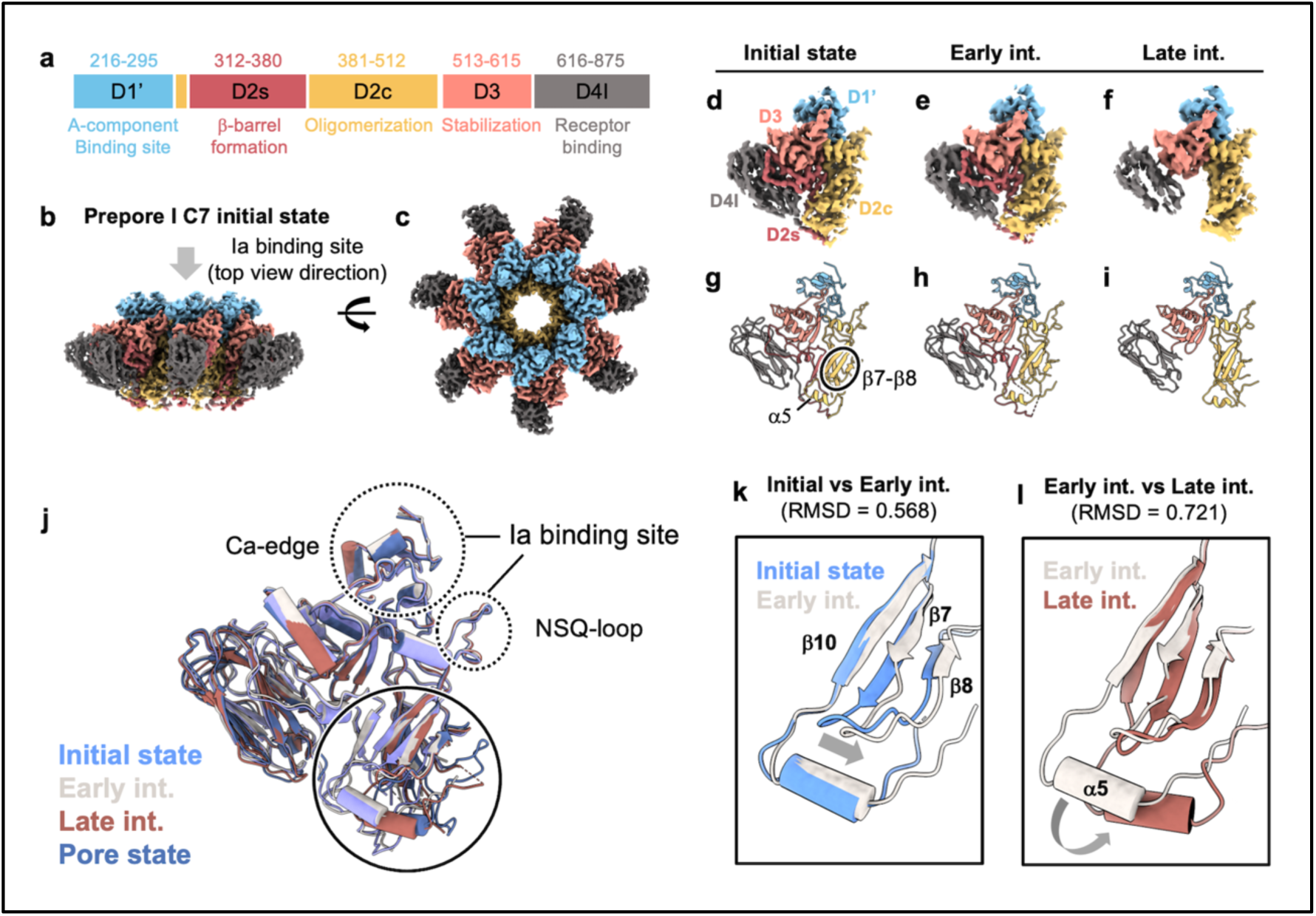
Structures of Ib protomer. **a** Schematic of the N-terminal cleaved Ib domains with densities observed in cryo-EM maps. **b, c** Cryo-EM maps of Prepore I, coloured according to the domains shown in panel **a**. **d−i** Maps and atomic models of the Ib protomer, coloured according to the domains. The Initial state map was extracted from the Prepore I, Early int. from Prepore II, and the Late int. in Prepore VIII. **j** Atomic models of the Initial state from Prepore I, Early int. from Prepore II, Late int. from Prepore VIII, and Pore state from Pore were aligned. **k, l** Comparison of cartoon models with tube α-helix between the Initial state (cyan) vs. Early int. (white), and Early int. vs. Late int. (brown). The RMSD values were calculated using UCSF Chimerax^45^.

During the single particle analysis of the Ib-pore, multiple pore states with varying degrees of β-barrel maturation were observed. These states exhibit no distinguishing features other than the length of the β-barrel. Focusing on the particle set that formed a fully extended β-barrel, particle images were selected for density map calculation **(Fig. 1k)**. Atomic models were refined against the density maps of the prepores and the pore, allowing for a comparative analysis of their structural characteristics.

### Structure comparison of Ib protomers

Ib consists of four domains: D1, which binds to the enzymatic component Ia; D2, which forms the pore lumen; D3, which interacts with other domains within a protomer and contributes to structural stabilisation directly; and D4, which is responsible for binding to the target cell receptor **(Fig. 2a-c)**. D2 can be further subdivided into D2c which serves as the structural foundation of the oligomer, and D2s which form β-barrel upon pore formation. Structural changes were observed in both of D2c and D2s. The constriction which was found in classification of Prepores (**Extended Data Fig. 2a),** corresponds to the conformational change of D2c toward centre of the oligomer. Based on the density of D2s, the protomer’s state could be roughly distinguished into two state as D2s visible protomer and D2s invisible protomer, which contributed as a key indicator to classify multiple prepore classes during the single particle analysis **(Extended Data Fig. 2c)**. In addition, atomic model building against the density map of D2c revealed that D2s visible protomer could be further subdivided into two classes due to the slight conformational difference, which we termed as Initial state **(Fig. 2d, g**) and Early int. **(Fig. 2e, h**), respectively. We also termed the D2s invisible protomer as Late int. **(Fig. 2f, i).** Consequently, the prepores comprised protomers in three distinct states, except for the pore state **(Fig. 2d-i, Supplementary Movie 1)**. Including the pore state, the most significant structural differences were observed for D2c and D2s **(Fig. 2j),** and no changes were observed in the Ca-edge and NSQ-loop which are the binding sites for enzymatic component Ia in each structure.

Among three protomer’s state, initial state was termed as it because the structure was the most similar to monomeric structural state of CDTb (PDBID:8DCM) and PA (PDBID:1ACC) reported in previous studies **(Extended Data Fig. 4)**^28,29^. On the other hand, both Early int. and Late int. had structural difference from initial state, which were partially similar to the Pore state. Compared to the initial state, the Early int. had a structure where β7 and β8 on the D2c domain at the protomer interface were shifted towards the CCW (counter-clockwise) **(Fig. 2k, Supplementary Movie 2)**, when top is defined as enzymatic component binding site and CCW is relative this orientation. In addition to the difference in D2s visibility, in the Late int., the position of α5 on the D2c domain was observed to shift towards the centre of the oligomer **(Fig. 2l)** comparing to the Early int. The Late int. was quite similar to the structure of the pore (Pore state), except that the β-barrel has not yet formed, and there was difference in the position of α5. A structural comparison of the Late int. and the Pore state is described in the following section. Based on these structural difference in D2s and D2c, the series of structures was suggested to change in the order of Initial state, Early int., Late int., and Pore state **(Supplementary Movie 1)**. The increase of backbone root mean square deviation (RMSD) value for Initial state compared to Early int. (0.568 Å), Late int. (0.789 Å), and Pore state (0.822 Å), supported this structural transition manner. Furthermore, the RMSD values for Late int. compared to Ealry int. (0.721 Å) and to Pore state (0.656 Å) suggested that Late int. has greater similarity with Pore state than Early int.

The arrangements of the three types of protomers that constituted the prepores obtained in this analysis were distinctive. From Prepore I to VIII, the number of protomers in the Initial state decreased by one, whereas the number of protomers in the Late int. increased by one. In addition, Late int. protomers were not scattered throughout the oligomer but were adjacent to each other **(Fig. 1a-h, Extended Data Fig. 5)**. These structural features suggested that the maturation of Pore completes through step-by-step conformational change from Prepore I to Prepore VIII in a domino-like manner, temporarily losing its symmetry. Another notable feature is that there was always one Early int. in the CW direction from a Late int.

### Conformational change at the protomers interface

It was necessary to consider what influences the adjacent protomer’s structure, which induces the conformational changes. Therefore, structural comparisons were made in pairs of protomers. In the Initial state-Initial state interface, the structure around β7-8 showed a gap **(Fig. 3a, b)**. In contrast, at the Early int.-Late int. interface, the structure was tightly zipped **(Fig. 3c, d)** by the shift of β7 and β8 which is observed in transition from Initial state to Early int. **(Fig. 2k)**, increasing the varied surface area of the interface **(Supplementary Movie 2)**. In other words, the initial structural change between the protomers is a process of reducing the solvent-exposed surface area and gaining interactions between the protomer pairs.

**Figure 3.**
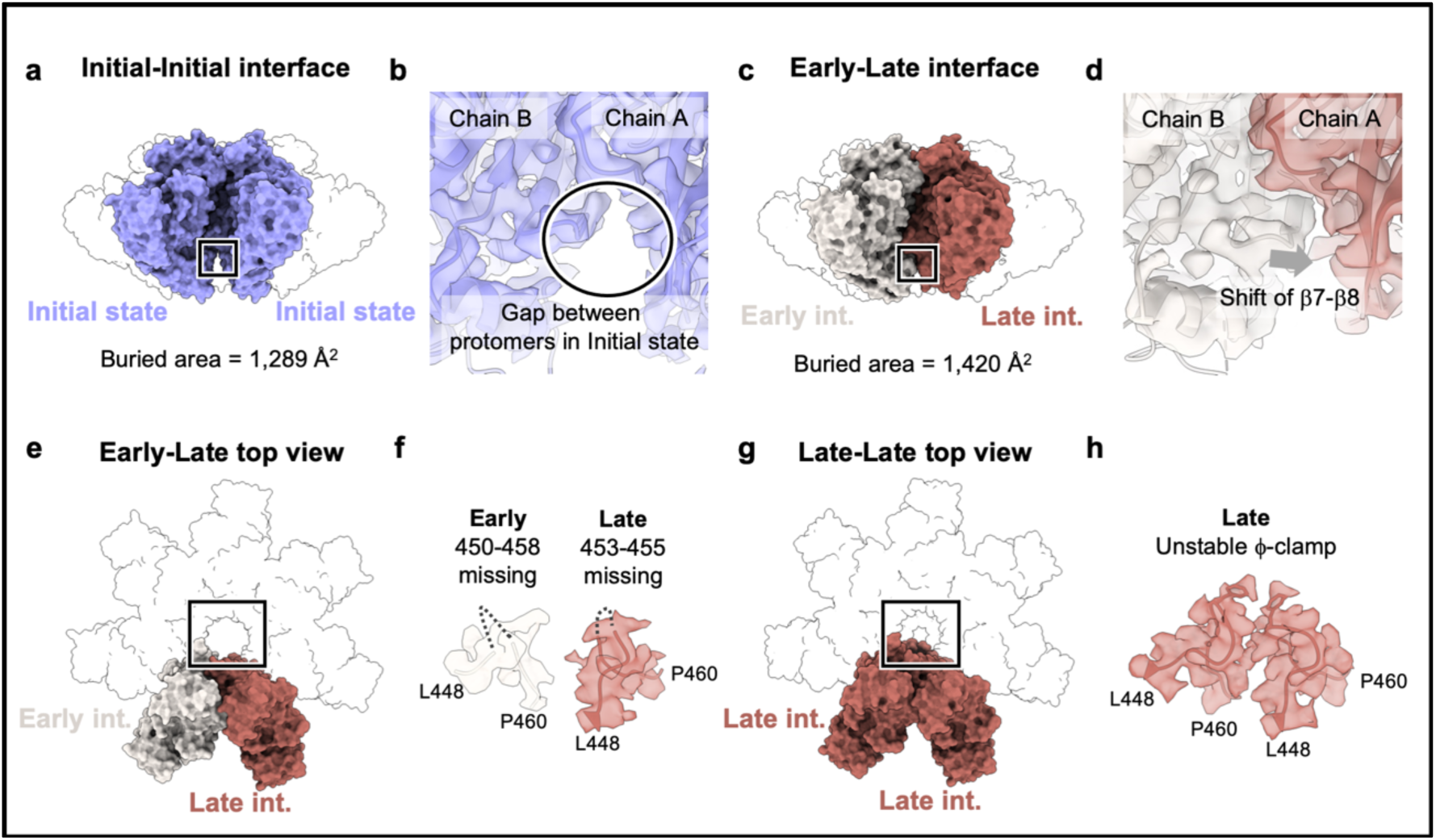
Comparison of inter-protomer interactions. **a−d** Comparison of interface structures between the Initial state and the Initial state (Prepore I) and the interface structure between the Early int. and Late int. (Prepore II). **a, c** Surface models of interest calculated from atomic models of Prepore I and Prepore II. **b, d** Atomic models in respective cryo-EM density maps. **e−h** Comparison of constriction-site structures between the Early int. and Late int. (Prepore II) and the interface structure between the Late int. state and Late int. (Prepore VIII). **e, g** Surface models of interest calculated from the atomic models of Prepores II and VIII. **f, h** Diagram of the atomic model in the ϕ-clamp map, generated with residues 448–460, and the density map trimmed around the atoms.

Next, the structural difference in the Early int.-Late int. interface and Late int.-Late int. interface were examined. However, no significant structural changes were observed at these interfaces. Prepore VIII formed by seven Late int. protomers, was found to have a relatively constricted structure with a more densely filled central density than earlier prepores **(Fig. 3e-h, Extended Data Fig. 6)**. This density could be explained as the residues forming the constriction-site known as the ϕ-clamp. Since the residues forming the ϕ-clamp (aa. 451−458) are located near α5 (aa. 460−473), it is reasonable that the shift of α5 toward the centre of oligomer observed from Early int. to Late int. led the formation of the ϕ-clamp **(Fig. 2l, Supplementary Movie 1, 3)**. The increase of protomers in Late int. through Prepore I to Prepore VIII allows gradual formation of ϕ-clamp, which facilitate protomers in Late int. to be adjacent each other resulting in domino-like manner and step-by-step transition to Prepore VIII. Although the ϕ-clamp forming residues in Late int. in Prepore II−VIII shift toward the centre of oligomer to form ϕ-clamp as well as Prepore VIII, the corresponding density was not observed in these maps, suggesting the structure is flexible. In Prepore VIII, although the density around the ϕ-clamp allowed for tracing the main chain structure, the side chains were not clearly observed even at the high resolution.

### High-resolution structure of Ib-pore

In our previous study, we determined the structures of Ib-pores solubilized with LMNG and the Ia-bound Ib-pore complex. The map of Ib-pore without Ia was obtained only in an immature short β-barrel state, while the map of Ia-bound Ib-pore was obtained in both short β-barrel and fully formed β-barrel states. Due to the Ia bound on to the Ib-pore, applying C7 symmetry during map calculation for the fully formed β-barrel was not feasible, which limited the resolution. In this study, we successfully obtained a high-resolution map of the Ib-pore with a fully formed β-barrel, allowing clear density observation up to the tip of the β-barrel, compared with previous research **(Fig. 4a, b)**. The Ib-pore contains several distinct constriction-sites within its lumen **(Fig. 4c)**. As described in the previous section, the Ca-edge and the NSQ-loop which are involved in binding to the enzymatic component, did not undergo structural changes from the prepore to the pore (**Fig. 2c)**. The most significantly changed constriction-site was the ϕ-clamp which was not observed in Prepore I, showed relatively unclear density in Prepore VIII (**Fig. 3g, h)**, and was clearly observed in Ib-pore **(Fig. 4d, Extended Data Fig. 6)**.

**Figure 4.**
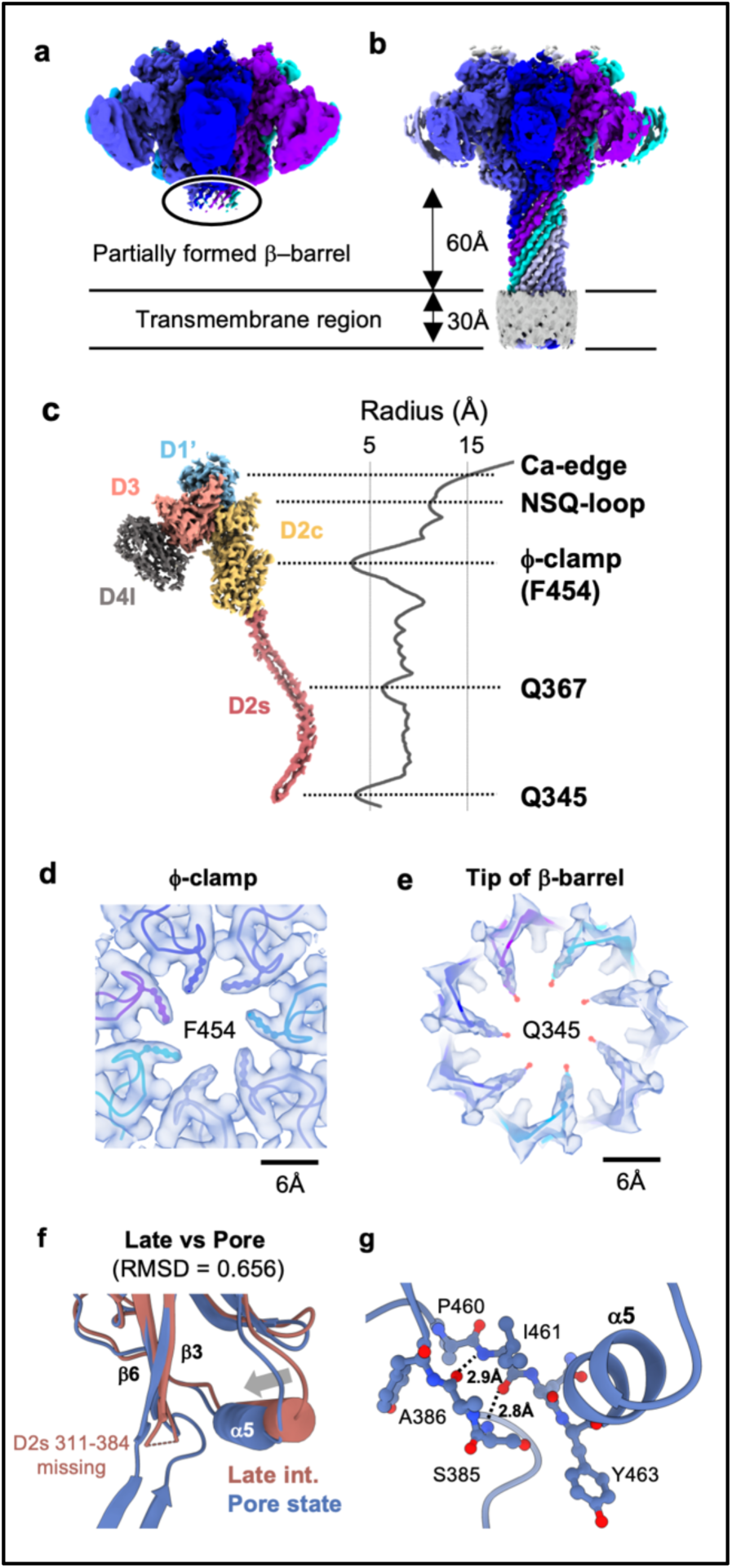
Structure of Ib-pore. **a, b** Cryo-EM maps of Ib-pores with immature-barrel (see Extended Data Fig.2a) and fully formed-barrel. The maps are coloured according to the protomers. **c** Cryo-EM map of an Ib protomer extracted from the Ib-pore with a fully formed barrel. The inner radius was calculated using HOLE2^49^ software. The positions of the constriction sites in the Ib-pore are marked with dotted lines. **d, e** Diagrams of the model showing the smallest inner radius. **f** Comparison of cartoons between Late int (brown) and the Pore state (purple). **g** Atomic model of Pore state Ib.

In the density obtained at the tip of the β-barrel, the hydroxyl groups of Y356 and Y344 were oriented outwards from the TM region, showing the characteristic snorkelling effect of transmembrane proteins. The amino acid side chains oriented inward in the β-barrel are rich in Ser/Thr and are relatively small. The side chain density of Q345, located at the hairpin turn of the β-barrel extended towards the centre, indicating a constriction **(Fig. 4c, e)**. However, the side chain density was relatively unclear and the side chains were far from each other, which suggested that the structure was flexible. Furthermore, in the Pore state, α5 was shown to shift towards the base of the β-barrel from Prepore VIII, forming interactions and stabilising the structure **(Fig. 4f, g, Supplementary Movie 3)**.

## Discussion

In this study, cryo-EM analysis of Ib-rosette which is the super-complex of a bacterial PFT revealed the step-by-step conformational changes as they transitioned from soluble prepore to membrane-spanning pore. Although the general mode of action of PFTs is widely recognised, detailed insights into their structural changes have mostly been based on comparisons of a few stable states and biochemical experiments with mutants. Being supported by cryo-EM density maps, we uncovered multiple structures including intermediate states during pore formation **(Fig. 1)**: the early immature state of soluble monomers forming a symmetric oligomer (Prepore I), asymmetric oligomers with structural changes in some protomers (Prepore II-VII), the fully matured symmetric Prepore VIII, and Pore with the membrane translocating the β-barrel. This is the first report for PFP family to reveal multiple intermediate structures in the membrane pore maturation process, supported by cryo-EM density maps without false symmetry by strategic analysis workflow (**Extended Data Fig. 2)**. These structural comparisons improve our understanding of conformational changes within individual protomers and allow for a more thorough discussion of the overall maturation process of binary toxin pores.

In our previous cryo-EM study of the Iota toxin, we determined the structures of both the Ib-pore and the Ia-bound Ib-pore complex (as well as the CDTa-bound CDTb-pore complex), revealing the docking mode of the enzymatic component as the initial step of its translocation through the Ib-pore, and showed that the N-terminus undergoes unfolding, which allows pass through ϕ-clamp which is the narrowest constriction-site of Ib-pore^20,21^. The results of this study revealed the detailed structures of the earlier steps, facilitated a more comprehensive understanding of the cellular entry mechanisms of binary toxins. Scott et al. also reported that the structural variety among PA protomers within the oligomer was classified into distinct states, using cryo-EM analysis of the D425A mutant of the anthrax toxin PA (which inhibits pore formation) based on the appearance of the density map of the receptor-binding domain. Moreover, protomers with similar structural states tend to be adjacent to each other within oligomer^26^. Expanding upon the previous report, we classified various protomer states based on their structural features on D2c and D2s using cryo-EM analysis and wild-type *Clostridium perfringens* iota toxin Ib samples. We identified the characteristic structural changes in the protomers and classified them into four states: Initial state, Early int., Late int., and Pore state. In addition, multiple structure of the entire oligomer differing in the ratio of each protomer’s state, were successfully identified. Furthermore, these protomer states were distributed in a distinctive pattern, in which the Late int. are always adjacent to each other at Early int. and was positioned in the most CCW direction within the oligomer, suggesting a regularity in the structural transition process **(Fig. 1a-h)**.

Initially, we aimed to prepare the Ib-pores in a lipid membrane environment using liposomes for solubilisation. However, the particles obtained were not proteoliposomes but rather aggregates of Ib oligomers assembled radially **(Extended Data Fig. 1a)**. Similar aggregates have been observed in the oligomers of the anthrax toxin PA, another binary toxin. It has been reported that such structures can be obtained by incubating oligomers bound to a carrier under acidic conditions^30^. However, unlike this study, structural analysis of the constituent oligomers has not been achieved. The antigen component of the influenza virus vaccine forms a radial structure composed of HA proteins referred to as the HA rosette^31,32^. Following this convention, we refer to the aggregated Ib structure as the Ib-rosette.

The Ib-rosette is an aggregate composed of multiple Ib oligomers; however, single particle structural analysis was possible by focusing on each oligomer. During the analysis, it was found that these oligomers included structures without β-barrels, partially formed β-barrels, and fully formed β-barrel **(Extended Data Fig. 1c-g)**. When observing the two-dimensional average images of the prepore class, an ambiguous density was found in the side view in the direction where the β-barrel would form **(Extended Data Fig. 1f, g)**. This indicated that other oligomers in Ib-rosette spatially hinder the formation of the β-barrel. In CDTb, it is known that paired heptamers form a dimer of heptamers; however, clear two-dimensional images of each heptamer are observed in these cases. In case of Ib-rosette sample, the average images of this prepore classes were clearly different^33,34^. We performed a more detailed classification of the prepore classes and obtained density maps for each of the eight prepores and the pore **(Fig. 1, Extended Data Fig. 2)**.

The multiple density maps obtained by Cryo-EM are independent of each other, making it difficult to interpret their correspondences. For example, in this study, it is not possible to establish a direct correspondence of protomers across different prepore states. Specifically, it remains uncertain whether the Early int. protomer observed in Prepore II undergoes a structural transition to the Late int. in Prepore III that is referred as CW model **(Fig. 5e-g)**, or retains its structure as an Early int. without any transition that is referred as CCW model **(Fig. 5h-j)**. At least, observations from a series of prepore maps suggest that the Ib-prepore adopts conformational changes through asymmetric intermediate structural states in step-by-step and domino-like manner. Based on these findings, we propose a possible structural transition models of the pore maturation in this study **(Fig. 5)**.

**Figure 5.**
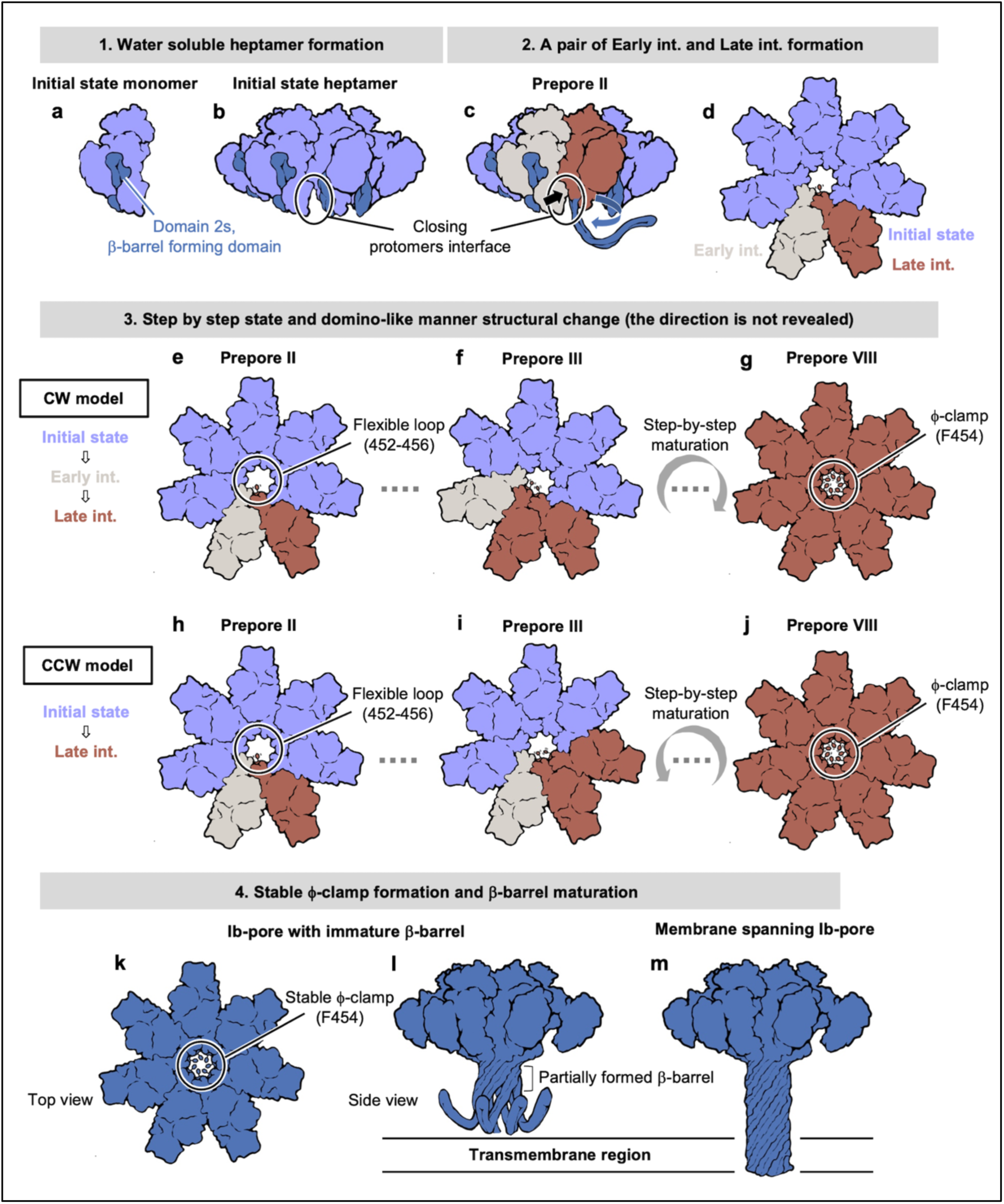
Summary of Ib-pore maturation mechanism. **a** N-terminal cleaved active Ib monomer in its Initial state. **b** Active Ib monomers assemble into heptameric Prepore I. **c** Two adjacent protomers convert into a pair of Early int. (white), and Late int. (Red) (Prepore II). **d** Top view of Prepore II. **e−g, h−j** Top views of the respective Ib-prepores. Number of Initial states and Late int. changed with step-by-step domino-like manner propagation in either CW or CCW direction. **g, j** The central constriction-site of the Ib-prepore is not yet fully formed but remains flexible. **k−m** Structures of the Ib-pores **k** Upon maturation to the pore, a stable ϕ-clamp is formed at the central constriction-site. The extension of the long β-barrel was achieved through the partial formation of the β-barrel from the foundation of the oligomer.

Comparison of the series of cryo-EM maps demonstrated formation of ϕ-clamp at the centre of the oligomer along with the pore maturation **(Extended Data Fig. 6)**. Phenylalanine residue forming ϕ-clamp is highly conserved among binary toxin family, and mutational study using anthrax toxin PA have revealed its critical role in maturation from the prepore to the pore^24^. The F454 residue of Ib is located upstream of α5 (aa. 460−473), which undergoes the most significant structural shift during the transition from Early int. to Late int. The density of the residues surrounding F454 (aa. 449−460) was not observed in the Initial state or Early int., suggesting that these residues possess a flexible structure in the states **(Fig. 3f)**. In the Late int., the density became more apparent especially in Prepore VIII, where the main-chain structure could be roughly traced. This observation indicates that the constricted region could be formed between a pair of protomers in Late int **(Fig. 3h)**. Because no significant structural changes were observed in the contact interface between the Late int.-Late int. pair and the pair that could be considered its precursor, we could not identify the factors contributing to the proximity of Late states. Although the most notable change in the Late int. is the movement of α5, this movement occurs away from the adjacent protomer, making it unlikely to stabilize the adjacent protomer into Late int. On the other hand, residues around the ϕ-clamp which show significant changes, are positioned to interact with each other. Therefore, the partial formation of the ϕ-clamp which is associated with the movement of α5, might facilitate to the tendency of Late int. states to come into proximity. Notably, despite of the high-resolution map of Prepore VIII at 2.9 Å, the side chains of residues forming ϕ-clamp was not observed, indicating the structural flexibility. Even at the similar resolution at 2.9 Å, the density is clearly observed in previously reported Ib-pore map (EMD-0721). Considering further structural shift of α5 during the transition from prepore VIII to pore as well **(Supplementary Movie 3)**, additional stabilization could be occurred, resulting in formation of stable ϕ-clamp in pore **(Fig. 5j, k)**.

The enzymatic components believed to pass through the pores exhibit different stoichiometries and binding positions in the iota toxin and anthrax toxins. In anthrax toxin, three enzyme molecules LF and EF bind to a single pore, with their N-terminal α-helices interacting with the α-clamp formed at the interface of PA protomers^35–37^, which makes the N-terminal α-helices face forwards to ϕ-clamp. In contrast, the single enzymatic components of the iota toxin and CDT bind to the Ca-edge of the pore, with its N-terminal helix unfolding at the NSQ-loop^20,21^. For binary toxin pores with a narrow constriction such as ϕ-clamp, the docking and unfolding of the enzymatic component from the Ca-edge to the NSQ-loop represent critical steps for activity. Notably, these regions remain unchanged in the structures of the individual monomer’s states **(Fig. 2j)**. This suggests that the enzymatic component could bind to the oligomer in both immature prepore state and mature pore states in same docking mode. In addition, the high-resolution cryo-EM density of the pore structure reported in this study revealed a constriction formed by Q345 at the tip of the β-barrel, which served as the exit for membrane translocation **(Fig. 4c, e)**. The diameter of this constriction site is less than 10 Å, which is smaller than the general diameter of an α-helix. Although a clear density was observed in the ϕ-clamp, the density of this constriction was ambiguous, suggesting that its structure is relatively flexible.

This study has enabled the discussion of the maturation process from Prepore to Pore in binary toxins at the molecular structural level, supported by the cryo-EM density **(Fig. 5)**. However, it should be noted that the observation of these structural states may have been made possible by the formation of the Ib-rosette, which potentially sterically hindered pore formation. Therefore, it is necessary to consider the possibility that the obtained structures may be artefacts, similar to samples in which pore maturation was cancelled by introducing mutations. However, the formation of the Ib-rosette may have enabled the trapping of unstable intermediates that also occurred during the normal transition from the prepore to the pore. Based on a comparison of the obtained structures, we proposed models of pore maturation with step-by-step process. Jiao *et al.* concluded that perforin undergoes a CW structural change by comparing temporal observations of the same molecule using atomic force microscopy (AFM) and specific directional structural perturbations through MD simulations^38^. During the maturation of the prepore, the number of Late ints. protomers increased sequentially. However, due to difficulty in identifying the correspondence of protomers in cryo-EM maps, whether this structural change proceeds via Early int. or directly from the Initial state, corresponding to the CW and CCW models respectively, remains to be investigated. In addition, the domino-like structural change in the prepore progressed through forming a pair of Early int. and Late int. as observed during the transition from Prepore I to Prepore II. However, it is also possible that the maturation process proceeds from more than two initiation points or occurs before heptamer formation. This hypothesis needs to be discussed, considering the speed of the maturation process and the likelihood of the emergence of additional initiation points. Future experiments involving single-molecule observations or simulations are expected to address these challenges and elucidate the detailed mechanisms of pore maturation.

PFPs are major components of bacterial toxins and the immune system and have become intriguing targets in both fundamental research and applications, such as their use as nanodevices in nanopore sequencing. However, these complicated proteins are produced as soluble monomers that mature into functional pores through dynamic structural changes. Binary toxins possess unique characteristics among pore-forming toxins that enable the translocation of enzymatic components into target cells via pores, making them highly applicable. Research is being conducted to create carriers by conjugating proteins to the binding domain of the enzymatic component, facilitating the intracellular delivery of the desired molecules^24,39^. This study is the first report of multiple structural states during the pore-forming process of a binary toxin through detailed classification using cryo-EM, which was previously difficult to obtain. Furthermore, we also expect that the findings of this study will facilitate our understanding of the general maturation processes of PFPs in general and contribute to the advancement of nanodevice development based on this structural foundation.

## Methods

### Expression and purification of Ib

Ib was expressed with a 20 kDa pro-sequence at the N-terminus which inhibits oligomer formation, and a 6-His tag at the C-terminus. Specifically, the gene encoding Ib (Uniprot ID: Q46221, residues 40−875) was cloned into pET23a without the signal sequence or other tags. The plasmid was transformed into *Escherichia coli* BL21 (DE3) and cultured in LB medium containing 50 ng/μl ampicillin at 37°C. When the culture reached an OD600 of 0.6, IPTG was added to a final concentration of 1 mM, and the culture was incubated for an additional 16 hours at 23°C. The cells were harvested by centrifugation.

The harvested cells were resuspended in buffer A (20 mM Tris pH 8.0, 100 mM NaCl, 1.0 mM CaCl2, 20 mM imidazole) containing one complete ULTRA Tablet (Roche, Basel, Switzerland) and lysed by sonication on ice. The lysate was subjected to ultracentrifugation (180,000*g*, 40 min) and the soluble fraction was bound to a Ni-NTA column. The column was washed with 10 volumes of buffer A and eluted with buffer B (20 mM Tris, pH 8.0, 100 mM NaCl, 1.0 mM CaCl2, and 500 mM imidazole). The eluate was concentrated by centrifugation using a 50 K ultrafiltration membrane and exchanged with buffer C (20 mM Tris, pH 8.0, 100 mM NaCl, 1.0 mM CaCl2).

### Sample preparation of Ib-rosette

Ib was activated and oligomerised in our previous studies^6^. Monomeric Ib was incubated at room temperature for 1 h with chymotrypsin at a 1/1,000 mass ratio. The reaction was stopped by adding PMSF (phenylmethylsulfonyl fluoride) at a final concentration of 1 mM to inhibit chymotrypsin activity. Ethanol was then added to a final concentration of 10% (v/v), and the mixture was incubated at 37°C for 1 h to promote Ib oligomerisation. 450 μl of the Ib oligomer solution was added to a Ni-NTA resin and washed with buffer C to remove ethanol. 540 μl of prepared liposomes (made by drying E. coli Extract Polar (Sigma-Aldrich) under N2 gas, resuspending with Buffer C, and processing through an extruder (Sigma-Aldrich) with a 100 nm filter), 360 μl of buffer C, and 100 μl of 1 M MES (pH 5.5) were then added, followed by incubation at 4°C for 16 hours. The resin was washed with ten column volumes of buffer A and eluted with buffer B. The eluted product was subjected to gel filtration using HiLoadTM 16/200 SuperdexTM 200pg of buffer C at a flow rate of 0.8 mg/ml. Fractions containing the Ib oligomer bands were collected based on SDS-PAGE observation and concentrated to 1.0 mg/ml using a 50 K ultrafiltration membrane.

### Data collection with Cryo-EM

The prepared sample (3 μl) was applied to hydrophilised Quantifoil R1.2/1.3, 300 mesh (Cu), and blotted for 7 seconds using a Vitrobot Mark IV (Thermo Fisher Scientific), followed by rapid freezing in liquid ethane. The frozen grids were imaged using Glacios (Thermo Fisher Scientific) equipped with a Falcon4 direct electron detector at the Institute for Life and Medical Sciences at Kyoto University to screen the grid preparation conditions through single-particle analysis. The final data were collected using a Titan Krios (Thermo Fisher Scientific) equipped with a K3 direct electron detector (Gatan), a BioQuantum energy filter (Gatan) with a slit width of 20 eV, and a Cs corrector (CEOS, GmbH) at the Institute for Protein Research, Osaka University, with a pixel size of 0.88 Å/pix, capturing 11,549 micrographs. The detailed imaging conditions are listed in Table 1.

**Table 1.** Statistics of cryo-EM data collection and single particle analysis.

| EMDB ID | Pore<br>EMD-61772 | Prepore I<br>EMD-61771 | Prepore II<br>EMD-61774 | Prepore III<br>EMD-61773 | Prepore IV<br>EMD-61775 | Prepore V<br>EMD-61776 | Prepore VI<br>EMD-61777 | Prepore VII<br>EMD-61778 | Prepore VIII<br>EMD-61779 |
| --- | --- | --- | --- | --- | --- | --- | --- | --- | --- |
| Magnification | 81,000 |  |  |  |  |  |  |  |  |
| Voltage (kV) | 300 |  |  |  |  |  |  |  |  |
| Electron Exposure (e/Å²) | 50 |  |  |  |  |  |  |  |  |
| Defocus Range (µm) | -0.6–1.6 |  |  |  |  |  |  |  |  |
| Pixel Size (Å) | 0.88 |  |  |  |  |  |  |  |  |
| Symmetry Imposed | C7 | C7 | C1 | C1 | C1 | C1 | C1 | C1 | C7 |
| Initial Particle Images (No.) | 6,223,563 |  |  |  |  |  |  |  |  |
| Final Particle Images (No.) | 161,249 | 10,511 | 15,199 | 22,352 | 8,878 | 11,285 | 7,676 | 23,305 | 17,097 |
| Map Resolution (Å) | 2.47 | 3.04 | 3.35 | 3.32 | 3.57 | 3.59 | 3.73 | 3.32 | 2.90 |
| FSC Threshold | 0.143 |  |  |  |  |  |  |  |  |

**Table 2.**
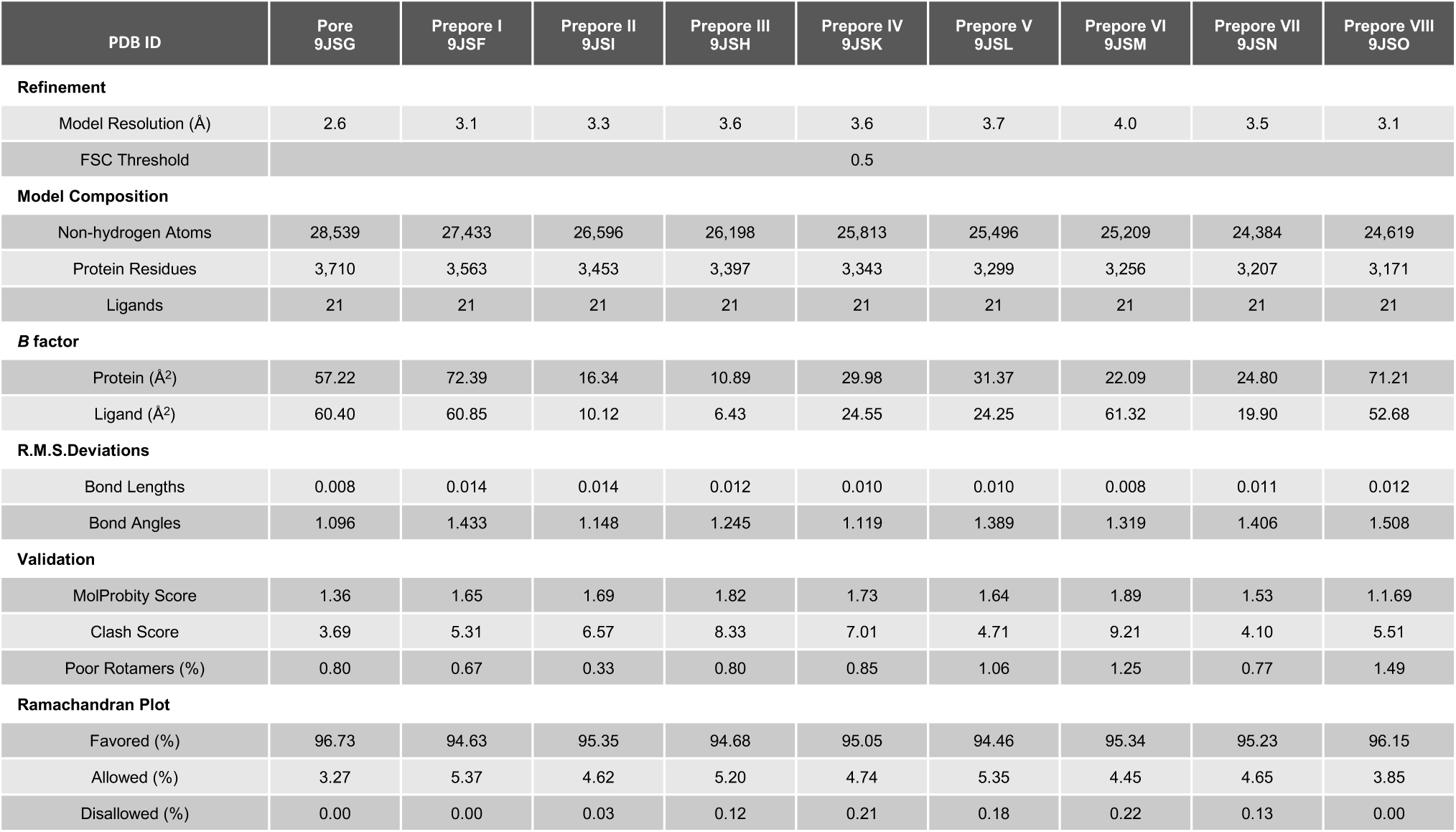
Statistics of atomic model refinement.

### Single particle analysis of Ib-pore

A series of analyses of the cryo-EM images was conducted using CryoSPARC (ver.3.3.2.)^27^. Motion correction was performed using Patch Motion Correction, and CTF estimation was performed using Patch CTF Estimation. Particle images were obtained using a Template Picker with a reference created by back-projecting a previously reported Ib-pore density map (EMD-0721). The particle images were extracted and downscaled to 1/4 of their original size. These extracted particles were roughly classified through Heterogeneous Refinement with C7 symmetry applied, resulting in a distinction between the prepore class, which lacks a β-barrel, and the pore class which has a β-barrel. The detailed analysis of the prepore class is described later in the “Classification of Ib-prepore” section. More homologous particle images were selected for the pore class through 3D Variability Analysis (3DVA)^40^ and no-align 3D classification. The resulting particle set was re-extracted without rescaling, and a 3D density map was reconstructed using Homogeneous Refinement. Additionally, Local Motion Correction^41^ was performed to correct for per-particle motion, followed by Homogeneous Refinement of images with defocus correction^42^. The local resolution was estimated with the RELION^43^ Local Resolution job **(Extended Data Fig. 3)**.

### Classification of Ib-prepore

3DVA was performed on the prepore class, resulting in the classification of circular oligomers into five distinct subclasses ranging from an expanded state to a more constricted state **(Extended Data Fig. 2a)**. Class 5 which is the most constricted class, exhibited a structure partially forming a β-barrel, similar to the Pore state; thus, further analysis was not conducted for this class. The remaining classes 1−4 were subjected to Symmetry Expansion with C7 symmetry, where the particle images were symmetrically replicated seven times. A mask was created focusing on one protomer, and the 3DVA of the protomer was used to distinguish between classes with and without an observable density corresponding to the D2s within the protomer **(Extended Data Fig. 2b, c)**. For Classes 1 and 2, the maps were reconstructed using Homogeneous Reconstruction Only with the particle sets where D2s were not visible, whereas Classes 3 and 4 used particle sets where D2s were visible. These reconstructions were references for refining the original particle sets before replication using Nonuniform Refinement. Among the resulting density maps, protomers where D2s were ambiguously visible, were subjected to focused non-aligned 3D classification to separate classes where D2s were visible from those where they were not; this process was iteratively repeated **(Extended Data Fig. 2e-g)**. Ultimately, the prepore states were classified into eight distinct classes, each differing in the number of protomers with clearly visible D2s. Finally, Non-uniform Refinement^44^ and Local refinement were performed for each of the eight prepore classes. The local resolution was estimated with the RELION^43^ Local Resolution job **(Extended Data Fig. 3)**.

### Model building

As structures such as the initial state and early int. where D2s are folded within the protomer, have not been previously reported in Ib, the predicted models were used as the initial model. The structure predicted from the Alphafold Database was edited to start the residue numbering from 1 after the signal sequence based on the amino acid sequence of Ib (Uniplot ID: Q46221), and used as the initial atomic model for the initial state. The predicted model was fitted as a rigid body into the most clearly observed initial state density in the Prepore I map using UCSF ChimeraX (ver.1.2.4)^45^, followed by manual model refinement using Coot (ver.0.9.5)^46^ and automated structural refinement using PHENIX (ver.1.19.2)^47^. For Early int., the refined initial state was used as the starting model. In contrast, for late int. and the pore state, the known Ib-pore structure (PDBID:6KLW) was used as the starting model and refinement was conducted against the respective density maps of Prepore II−VIII, and Pore in the same manner as described above. Although the D4-I structure of Ib has not been previously reported, its density was observed in Prepore I. Therefore, the refined D4-I structure from Prepore I was used as the initial model for other oligomers and structural refinement was conducted again using PHENIX. Finally, the chain orientations were validated using MolProbity^48^ for the models obtained using the above process. Inner radius of Ib-pore was calculated using HOLE2^49^ software.

## Data availability statement

The Cryo-EM maps and coordinates were deposited to the Electron Microscopy Data Bank (EMDB) and Protein Data Bank (PDB) with the accession codes EMD-61772 and PDBID 9JSG for Pore state, EMD-61771 and PDBID 9JSF for Prepore I, EMD-61774 and PDBID 9JSI for Prepore VII, EMD-61773 and PDBID 9JSH for Prepore III, EMD-61775 and PDBID 9JSK for Prepore IV, EMD-61776 and PDBID 9JSL for Prepore V, EMD-61777 and PDBID 9JSM for Prepore VI, EMD-61778 and PDBID 9JSN for Prepore VII, and EMD-61779 and PDBID 9JSO for Prepore VIII. The cryo-EM images are deposited to Electron Microscopy Public Image Analysis (EMPIAR) with the accession code EMPIAR-12333.

## Acknowledgements

This work was supported by JSPS KAKENHI Grant Numbers 21H02452 (to H.T.), 24K01993 (to H.T.), and 21J13410 (to T.Y). This research was partially supported by the Platform Project for Supporting Drug Discovery and Life Science Research (Basis for Supporting Innovative Drug Discovery and Life Science Research (BINDS)) from AMED under Grant Number 2366 and the Joint Usage/Research Center Program of the Institute for Life and Medical Sciences at Kyoto University (to H.Y., Y.S., and T.N.). We thank J. Kishikawa, M. Hirose, and T. Kato at the Institution for Protein Research, Osaka University, for training on the techniques used for the cryo-EM study. We would like to thank Editage (www.editage.jp) for English language editing

## Author contribution

All author participated in research design; T.Yamada prepared the Ib-rosette sample for cryo-EM; T.Yamada, Y.S and T.N performed cryo-EM data collection, and T.Yamada and Y.S performed single particle analysis; T.Yamada and T.Yoshida performed the atomic model building; all authors contributed to the discussion and writing the manuscript, and H.T supervised the project.

## Author information

Competing interests: The authors declare no competing interests.

---

**Extended Data Figure 1.**
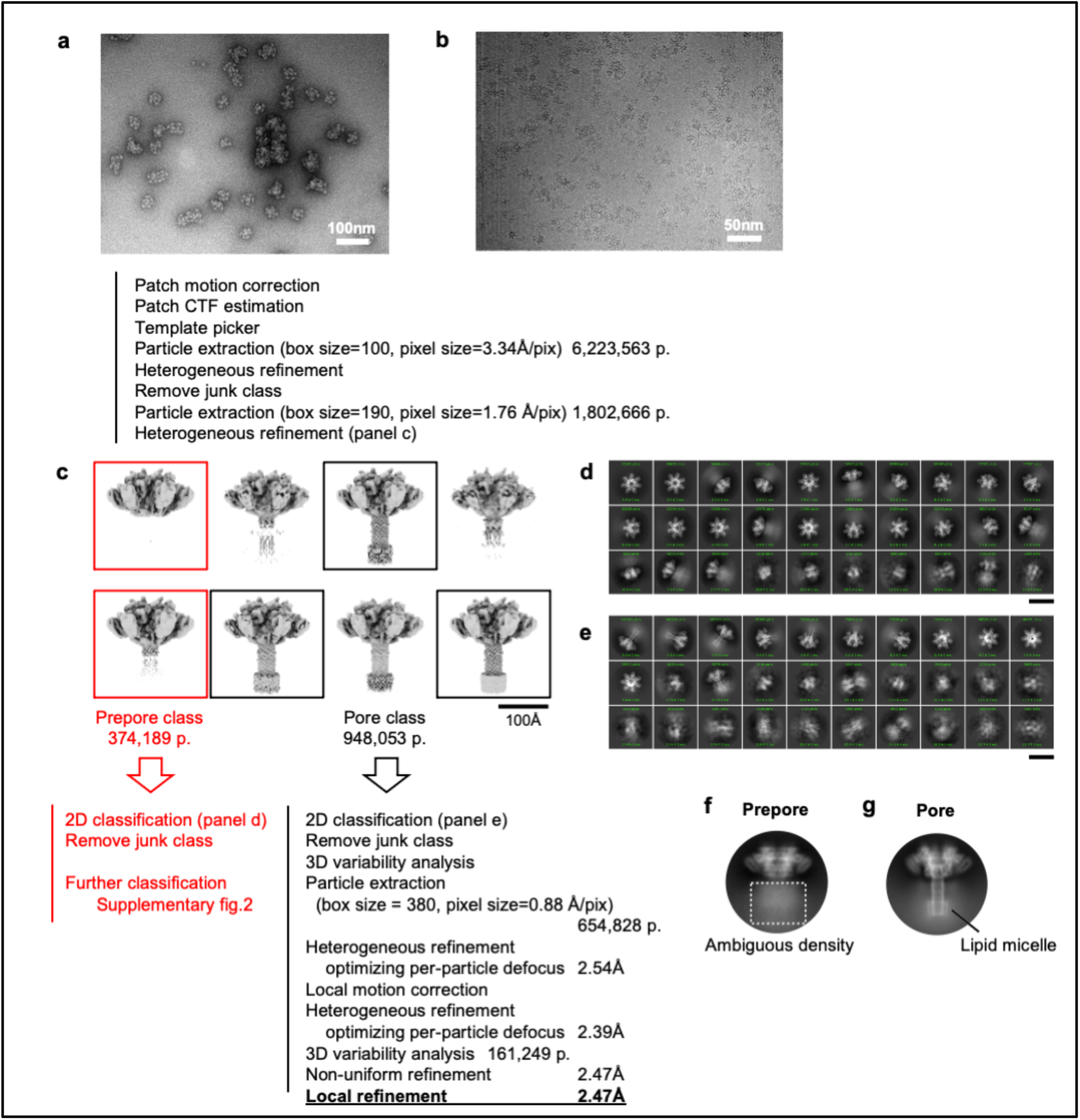
Single particle analysis workflow. **a** Micrograph of negatively stained Ib-rosette. **b** Cryo-EM micrograph of the Ib-rosette. **c** Representative density maps calculated using heterogeneous refinement. The classes marked with squares were manually sorted into the Ib-prepore particle set (red) and Ib-pore particle set (black). **d, e** Averaged images of Ib-prepores and Ib-pores calculated using 2D classification. The scale bar represents a length of 200 Å. **f, g** Representative 2D average images of the side views of the Ib-prepores and Ib-pores.

**Extended Data Figure 2.**
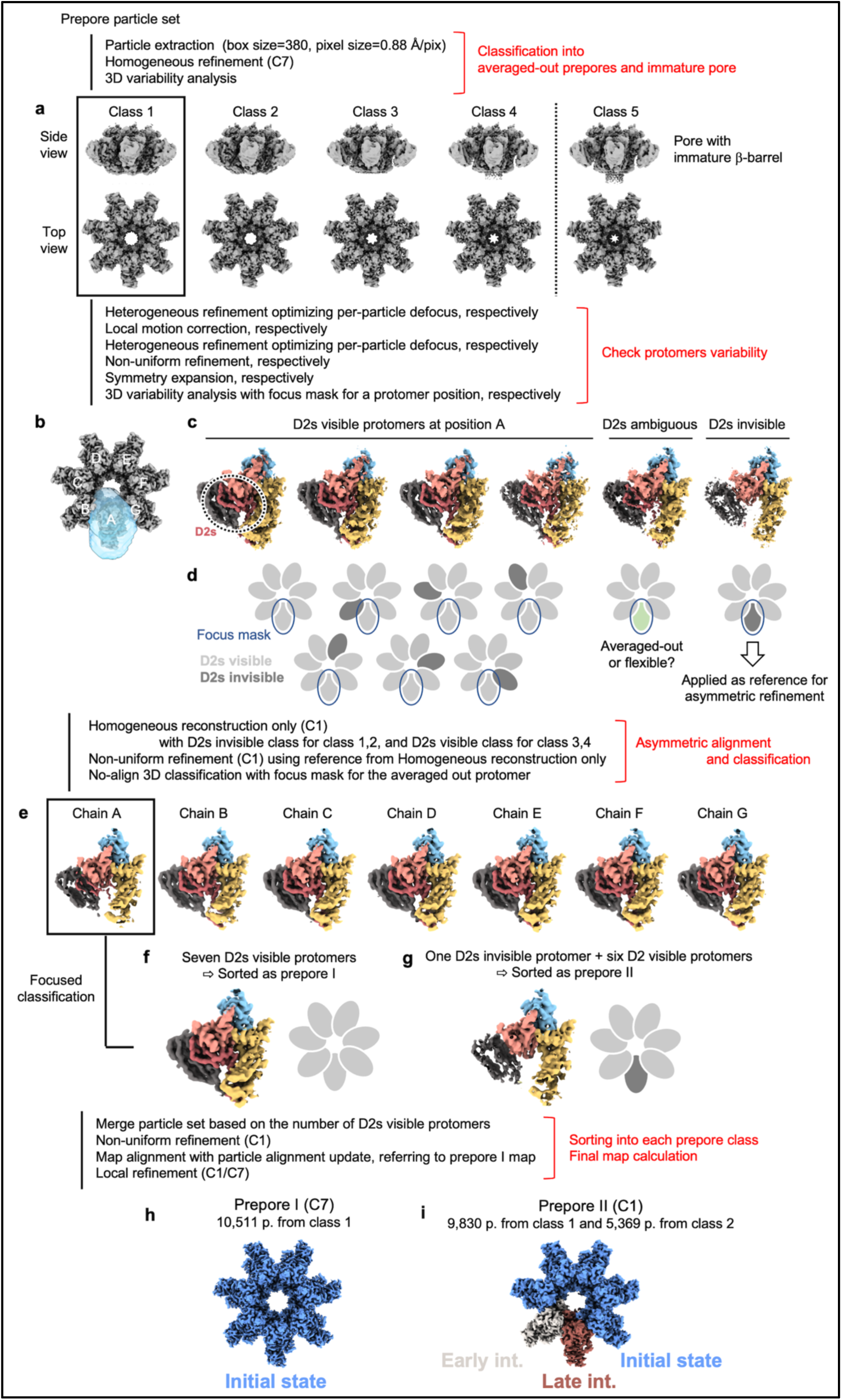
Detailed classification workflow for Ib-prepore. **a** Classes obtained from 3D variability analysis. **b** Density of subunit A (transparent light blue) indicates the mask used for focused classification of symmetry-expanded particles. **c, d** Classes obtained from focused 3D variability analysis and the corresponding diagram of the possible relative positions between the focused mask and D2s-visible or D2s-invisible protomers. **e** Protomer maps extracted from the non-uniform refinement output. Subunit A shows a weak D2s density because the D2s-visible protomer (**f**) and the D2s-invisible protomer (**g**) are aligned at the same position. **h, i** Ib-prepore maps obtained from Class 1 particles.

**Extended Data Figure 3.**
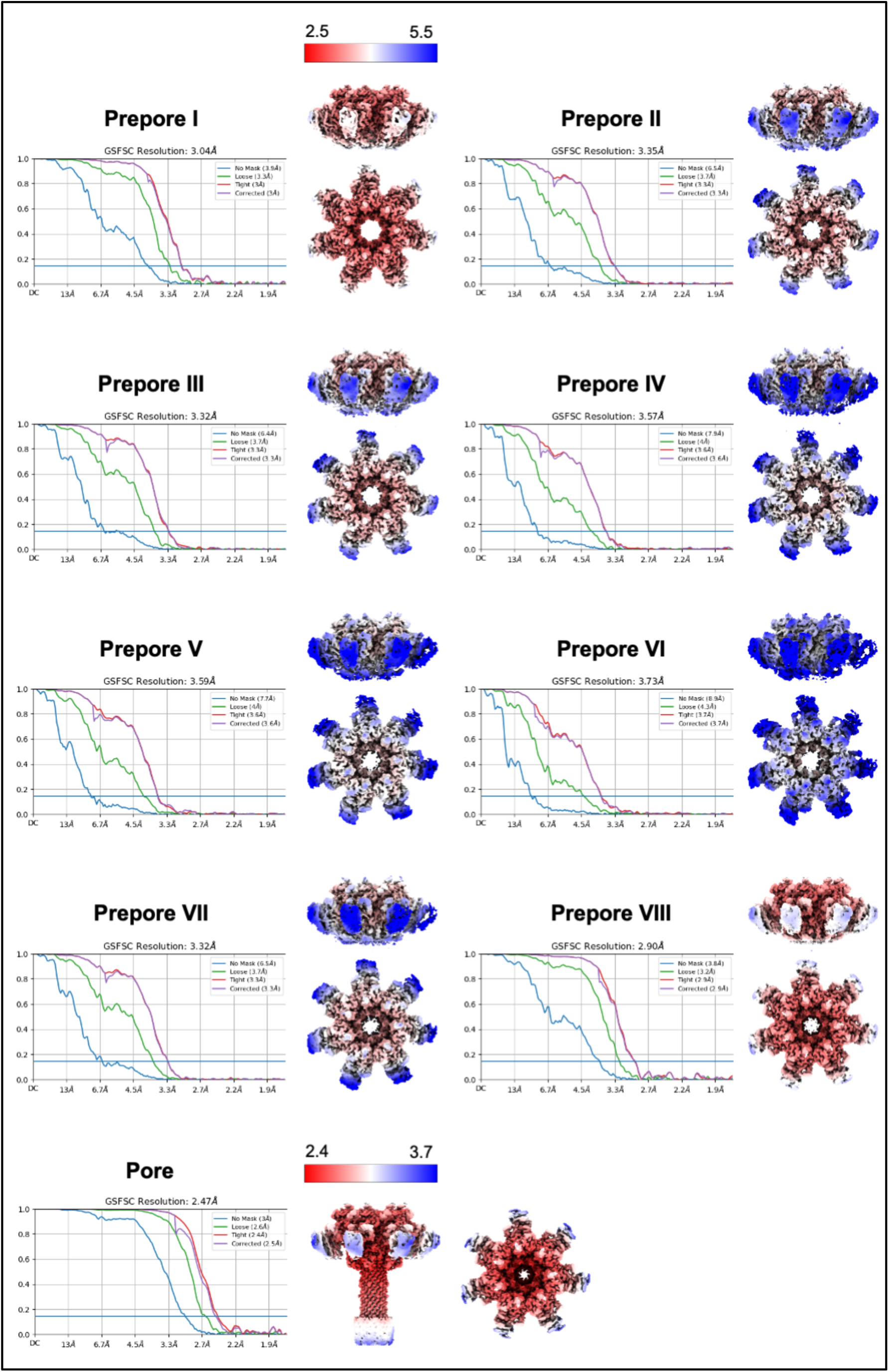
Global and local resolution. The global resolution was calculated using the gold-standard FSC threshold of 0.143. The local resolution was estimated with the RELION Local Resolution job using the final half-maps that are calculated using CryoSPARC. For Ib-prepore, the maps are coloured by the same resolution range (2.5 Å to 5.5 Å), while an independent resolution range coloured Ib-pore due to its outstanding resolution.

**Extended Data Figure 4.**
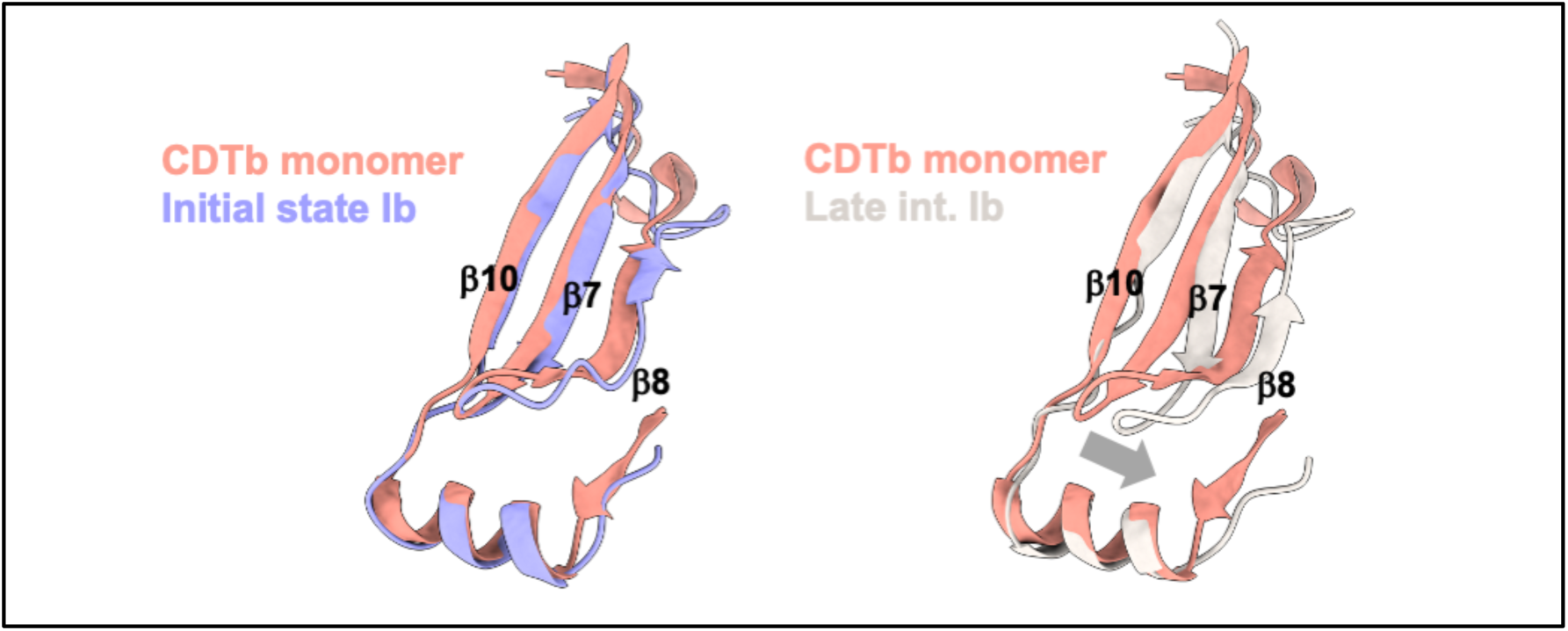
Structural comparison between Ib and CDTb monomer. The monomeric CDTb structure (PDBID: 8DCM) was aligned to the Ib Initial state from Prepore I and the Early int. from Prepore II.

**Extended Data Figure 5.**
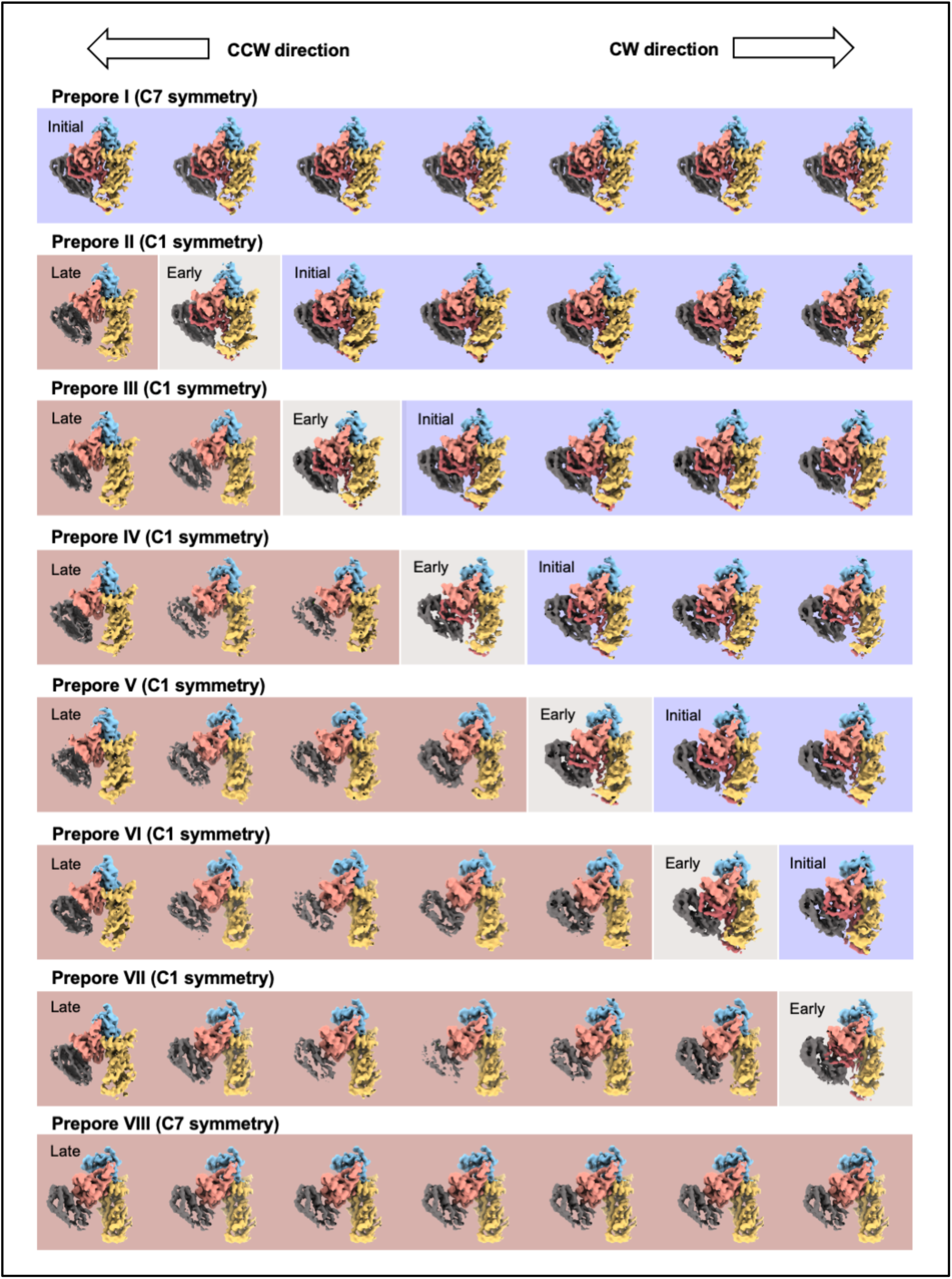
Protomers profile contained in oligomers. Density maps of the protomers were trimmed from their respective prepores, along with their atomic models. Owing to the symmetry for averaging, applied during the single particle analysis, all protomer maps from Prepore I and Prepore VIII are same. The CW and CCW directions were defined relative to the top of the Ib oligomer where the Ia docking region was located.

**Extended Data Figure 6.**
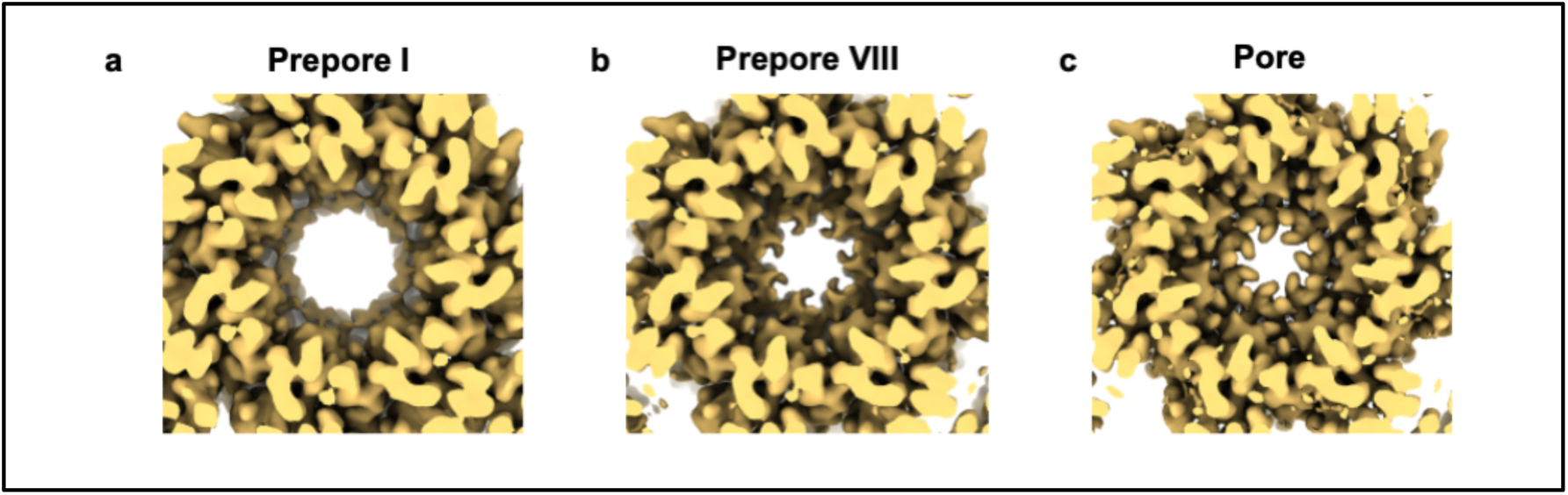
Comparison of the constriction-site top view. A close-up view of the central constriction site. Each map was calculated using C7 symmetry.

**Supplementary Movie 1 | Comparison of monomer structures**

The atomic models of a protomer in Prepore I, Prepore II, Prepore VIII and the Pore are utilized to represent the Initial, Early int., Late int. and Pore states, respectively.

**Supplementary Movie 2 | Structures of interface between the protomers in Initial state−Initial state and protomers in Early int.−Late int.**

The movie was created using atomic models and cryo-EM density maps of Prepore I and Prepore II.

**Supplementary Movie 3 | Shift in the position of α5 during structural transitions**

The video was generated using atomic models arranged to reflect conformational changes propagating in the CW direction. Reverse propagation of the conformational changes in the CCW direction may occur as well.

